# Hexokinase 2 expression in apical enterocytes correlates with inflammation severity in patients with inflammatory bowel disease

**DOI:** 10.1101/2024.04.04.588060

**Authors:** Saskia Weber-Stiehl, Jan Taubenheim, Lea Järke, Christoph Röcken, Stefan Schreiber, Konrad Aden, Christoph Kaleta, Philip Rosenstiel, Felix Sommer

**Affiliations:** Institute of Clinical Molecular Biology, University of Kiel, Rosalind-Franklin-Straße 12, 24105 Kiel, Germany; Institute of Experimental Medicine, University of Kiel, Michaelisstr. 5, 24105 Kiel, Germany; Department of Pathology, University Hospital Schleswig-Holstein, Campus Kiel, Arnold-Heller-Straße 3/House U33, 24105 Kiel, Germany; Department of Internal Medicine I, University Hospital Schleswig-Holstein, Campus Kiel, 24105 Kiel, Germany

**Keywords:** inflammation, hexokinase, HK2, human biopsies

## Abstract

**Background:** Inflammation is characterized by a metabolic switch promoting glycolysis and lactate production. Hexokinases (HK) catalyze the first reaction of glycolysis and inhibition of epithelial HK2 protected from colitis in mice. HK2 expression has been described as elevated in patients with intestinal inflammation, however there is conflicting data from few cohorts especially with severely inflamed individuals, thus systematic studies linking disease activity with HK2 levels are needed.

**Methods:** We examined the relationship between HK2 expression and inflammation severity using bulk transcriptome data derived from the mucosa of thoroughly phenotyped patients suffering from intestinal inflammation of two independent cohorts. Analyzing publicly available single cell RNA sequencing data and performing immunofluorescence on colonic biopsies of unrelated patients with intestinal inflammation confirmed the RNA-based findings on cellular and protein level.

**Results:** HK2 expression gradually increased from mild to intermediate inflammation, yet strongly declined at high inflammation scores. Expression of epithelial marker genes also declined at high inflammation scores, whereas that of candidate immune marker genes increased, indicating a cellular remodeling of the mucosa during inflammation with an infiltration of HK2-negative immune cells and a loss of the apical epithelium – the main site of HK2 expression. Normalizing for the enterocyte loss clearly identified epithelial HK2 expression as gradually increasing with disease activity and remaining elevated at high inflammation scores. HK2 protein expression was mostly restricted to brush border enterocytes and these cells along with HK2 levels vanished with increasing disease severity.

**Conclusions:** Our findings clearly define dysregulated epithelial HK2 expression as an indicator of disease activity in intestinal inflammation and suggest targeted HK2-inhibition as a potential therapeutic avenue.

## Background

Prevalence of inflammatory bowel disease (IBD) is rising globally causing severe health issues and drastically reducing quality of life. These diseases are multifactorial with complex interactions of multiple genetic and environmental factors^1^. Despite decades of intense research, the exact etiology of IBD remains mostly unknown limiting treatment options^2^. In recent years it became evident that ongoing inflammation is characterized by changes in metabolic activity, in particular glucose metabolism^3–5^. Hexokinases (HK) catalyze the first and irreversible step of glycolysis – the phosphorylation of glucose to glucose-6-phosphate – and are therefore crucial for glucose metabolism and maintenance of homeostatic body functions. In mammals, five HK isoenzymes have been identified: HK1, HK2, HK3, GCK (Glucokinase) and HKDC1 (HK domain containing 1). Each of these HK isoenzymes display specific tissue expression patterns and glucose affinities. HK2 is of particular interest as it is the most abundant HK member within the intestine, exerts high HK activity, responds to various internal as well as external stimuli and shows elevated levels during inflammation^6^. Recently, we further revealed an upregulated expression of HK2 specifically in inflamed compared to non-inflamed tissue of the same patient, irrespective of the type of intestinal inflammation and demonstrated that ablation of HK2 in intestinal epithelial cells (IEC) protects from acute intestinal inflammation, suppresses cell death and alters mitochondrial function^6^. These findings placed HK2 as a molecular target to treat intestinal inflammation. However, when expanding our research about the role of HK2 in intestinal inflammation, we encountered transcriptome datasets with contradictory findings of either an unaltered HK2 expression or even downregulation, especially in severely inflamed patients undergoing bowel resection^7^. We therefore aimed to investigate the cause of this apparent contradiction with the goal to clarify the relation between HK2 expression and intestinal inflammation. Here, we show that with increasing inflammation severity the intestinal mucosa is gradually remodeled, which comprises a partial loss of the apical epithelium – the primary source of HK2 expression – and a simultaneous infiltration of immune cells. Normalizing for this cellular remodeling clearly demonstrates a gradual upregulation of HK2 expression with severity of intestinal inflammation.

## Methods

### Human studies

We used mucosal transcriptomic data and fixed intestinal specimen of two independent large longitudinal clinical studies, namely the EMED^9^ (191 samples) and FUTURE^8^ (87 samples) cohorts (trial IDs: EudraCT number 2016-000205-36 and ClinicalTrials.gov NCT02694588), for which accompanying disease activity scores (Harvey-Bradshaw Index (HBI)/Mayo Score) were available (see Table S1 for cohort clinical characteristics). Note that multiple samples were collected from each patient. Sample origin was incorporated in all following analyses. The HBI^26^ is used to quantify disease activity in patients suffering from Crohn’s disease (CD), one of the two IBD subtypes. Here, variables such as general well-being, abdominal pain and abdominal mass with each having scores of 0-3, as well as the number of liquid stools per day (1 point per stool) and other complications (1 point per complication) yield the “open-ended” total score (Table S3). The Mayo score is used for ulcerative colitis (UC), the other clinical IBD subtype. This scoring system accounts for general well-being, rectal bleeding, endoscopic results and stool frequency with scores of 0-3 per category for an overall score ranging from 0 to 12 (Table S3). To facilitate a direct comparison of both scores despite them being “open-ended” and “discrete”, we calculated a general inflammation score by setting the highest score in the dataset for each HBI and Mayo score to 1 and scaled the score of each patient accordingly (see Table S3).

### Transcriptome data analysis

RNA sequencing data derived from mucosal biopsies of patients with various grades of intestinal inflammation were retrieved from two previously published IBD cohorts^8,9^. Read counts were transformed to Transcripts Per Million (TPM) values to normalize for differential sequencing depths among samples. TPM data were then plotted per sample against the inflammation score using R (version 4.2.2) and ggplot2 (3.4.3). Trendlines were calculated including all data points and using the distance to the actual location as a weight to enable a robust calculation and avoiding overfitting especially in areas of scarce sampling. To test for an association of gene expression to changes in the inflammation score we used variance-stabilization transformation of raw read counts and fitted linear mixed models with following form: vst(geneExpression) ∼ Age + Sex + Diagnosis + InflammationScore. Statistical analyses were performed using R (version 4.2.2^27^ with the following packages: lm4 (version 1.1-29)^28^, lmerTest (version 3.1-3)^29^, car (version 3.0-13)^30^, ggplot2 (version 3.3.6)^31^ and DESeq2 (version 1.38.3)^32^.

### Cellular deconvolution

We deconvoluted the bulk RNA data using the MuSiC package (version 1.0.0)^10^ with default values. As reference, we used a single cell data set from UC patients described in a study by Smillie et. al.^11^. The original single cell data set was split in “epithelial”, “immunogenic” and “stromal” subsets, which we rejoined in our analysis. Due to limited resolution in deconvolution approaches and to reduce cell diversity, we pooled ontogenetically closely related cell types (Table S4). We restricted the analysis to the eight most abundant cell types.

### Single cell RNA sequencing

Single cell data was obtained from the Single Cell Portal (accession SCP259) initially described by Smillie and colleagues^11^. The different cell populations were rejoined and general quality controls were performed. In short, we filtered cells with read counts below 1000 (low quality). Further, we removed cells with less than 200 or with more than 2720 (two-fold of the standard deviation of expressed genes across all cells) differently expressed genes to account for spurious sequencing depth and removal of duplicates. All cells with a mitochondrial RNA content above 5% were also removed. For plotting, gene expression values where log-transformed. Plotting and quality filtering was performed in R (version 4.2.2) using ggplot2 (version 3.4.3) and Seurat (version 4.4.0)^33^.

### Immunofluorescence

5 µm sections of paraffin-embedded colonic biopsies were deparaffinized with Xylol substitute (Roth), incubated in citrate buffer for 3 minutes and subsequently blocked in 5 % BSA-PBS and 0,2 % TritonX for 30 minutes. Primary anti-E-cadherin (1:400 in 1 % BSA, #3195, Cell Signaling Technology) and anti-Hexokinase 2 (1:500 in 1% BSA, Cat# NBP2-02272, Novus Biologicals, Colorado, US) antibodies were incubated over-night. Sections were washed, incubated with secondary antibodies (Alexa Fluor 488 goat anti mouse, Invitrogen, A32731 and Alexa Fluor 555 goat anti rabbit, Invitrogen, A21430) for 45 minutes at room temperature. DAPI (1:40000 in PBS, D9542, Sigma Aldrich, St. Louis, US) was used for DNA counterstaining. Slides were mounted using antifade mounting media (DAKO, Hovedstaden, Denmark). The quantitative analysis was performed using fluorescence microscopy for stained samples using the imager Z1 microscope (ZEISS, Jena, Germany) and ZEN software (version 3.0). Images were taken by a digital camera system (AxioCam HrC/HrM, Zeiss, Jena, Germany) and ApoTome (ZEISS, Jena, Germany). Fluorescence signal intensity was measured using Fiji/ImageJ software.

### Statistics

General statistical analyses were performed using the GraphPad Prism 9 (GraphPad Software Inc., La Jolla, USA). For pairwise comparisons the Mann-Whitney-U test was used whereas for multiple comparisons one-way ANOVA with false discovery rate (FDR) correction were performed. Data are shown as mean ± standard error of the mean (SEM). A p-value of ≤ 0.05 was considered as significant (*). A p-value of ≤ 0.01 was considered as strongly significant (**) and p-value of ≤ 0.001 as highly significant (***). For longitudinal data, linear mixed effect models were fitted where gene expression was used as dependent variable, while inflammation score and patient ID were fixed and random factors. The models were fitted using lme4 (version 1.1-31) and statistical testing for coefficients was performed using lmerTest (version 3.1-3). Model validity was tested against a null-model using log-likelihood ratio tests and model assumption were evaluated by diagnostic plotting of model residuals, data point influence strength and random factor fitting.

## Results

### Epithelial HK2 expression increases with high inflammation scores

To determine a possible correlation of HK2 expression and disease activity, we analyzed RNA sequencing (RNA-seq) data derived from intestinal biopsies of patients suffering from IBD of two independent clinical studies^8,9^, for which accompanying disease activity scores for each sample were available (Table S1). This analysis revealed that HK2 expression initially increased gradually reaching peak expression at mid disease activity (inflammation score ∼ 0.4) and then declined with high inflammation scores (Figure 1A). As within the intestine HK2 is mainly expressed by epithelial cells of the apical mucosa (Figure S1), we hypothesized that with very high levels of inflammation these apical epithelial cells and therefore also the main site of HK2 expression might be lost due to shedding and apoptosis or transdifferentiation resulting in decreased HK2 expression levels in bulk RNA-seq data. To investigate this hypothesis, we assessed the expression levels of three different epithelial marker genes, namely E-cadherin (*ECAD*), epithelial cell adhesion molecule (*EPCAM*) and Villin1 (*VIL1*). Supporting our hypothesis, the expression levels of these epithelial markers were all declining with increasing inflammation severity (Figure 1B-D), which indicates progressive epithelial damage caused by the inflammation. To account for this epithelial loss, we then normalized the expression of HK2 to that of these epithelial markers, thereby yielding an epithelial HK2 expression. Importantly, epithelial HK2 expression gradually increased in correlation to disease activity and then remained elevated at high inflammation scores (Figure 1E). To test whether the observed expression changes were unique to the combined clinical cohorts, we also performed these analyses separately for each clinical cohort (Figure S2). Importantly, we observed for both clinical cohorts the same changes in *HK2*, *ECAD*, *EPCAM*, *ECAD* and epithelial HK2 expression. We also tested for potential differences between the two main types of IBD, Crohńs disease (CD) and ulcerative colitis (UC) and found similar responses both for CD and UC (Figure S3). Finally, correlation of HK2 expression with disease severity was confirmed using linear regression models (Figure S4 and Table S2).

**Figure 1:**
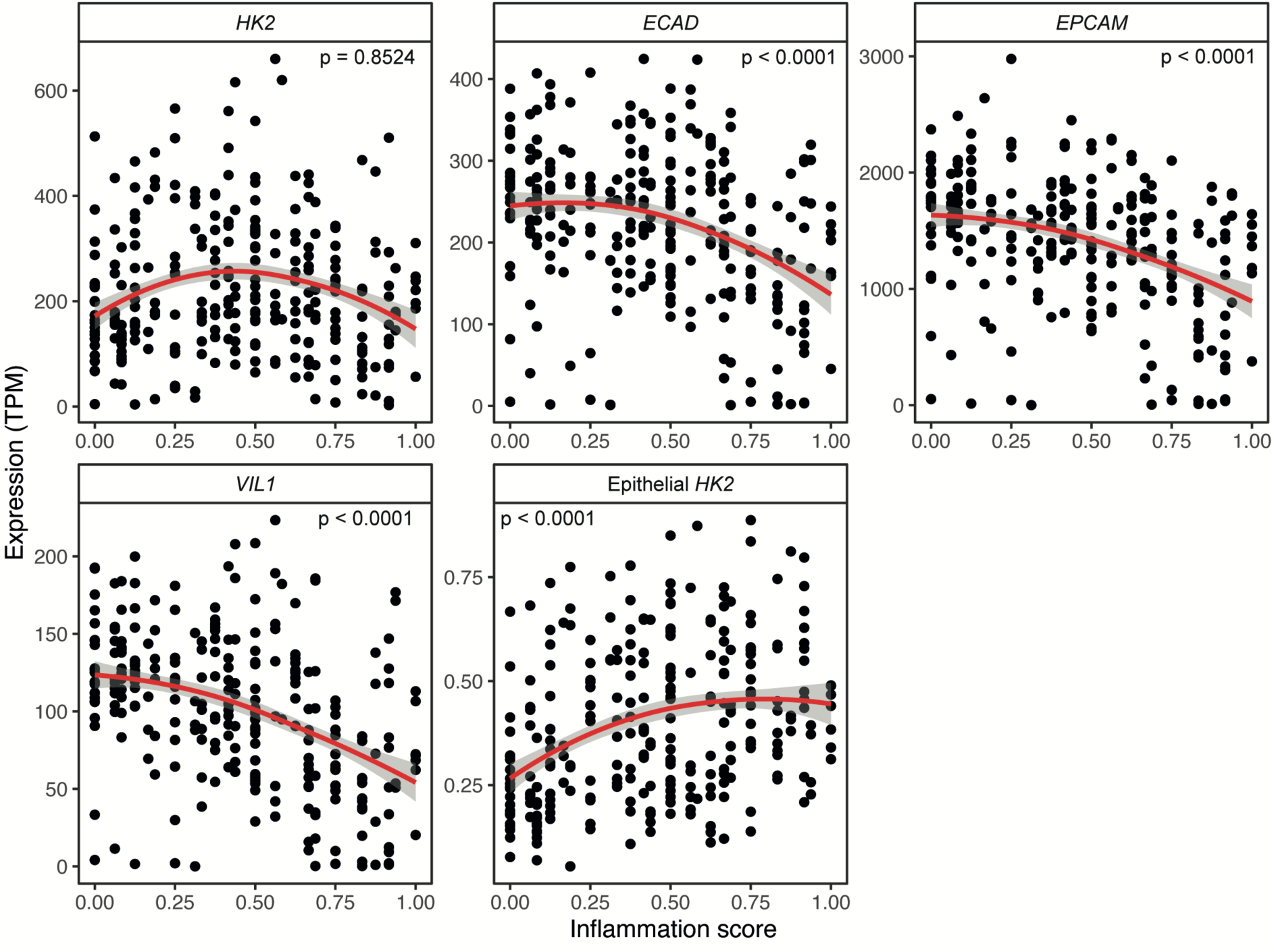
Epithelial HK2 expression correlates with inflammation severity. **A-D)** Expression (TPM, transcript per million) of *HK2* **(A)** and the epithelial marker genes *ECAD* **(B)***, VIL1* **(C)** and *EPCAM* **(D)** in the intestinal mucosa of patients with various degrees of gut inflammation. Note that at high inflammation scores (>0.4) expression of *HK2* and the epithelial marker genes all decrease indicating epithelial erosion. The inflammation score was calculated as a scaled Harvey-Bradshaw index (Crohn’s disease) or Mayo score (ulcerative colitis) to accommodate both disease types. **E)** *HK2* expression increases with inflammation scores after normalization to epithelial marker gene expression. The red lines represent the mean expression trendline. The number in the upper right corner represents the p value for the correlation between gene expression and inflammation score as determined by linear mixed model.

### Immune cell infiltration during intestinal inflammation

Another source for cellular remodeling during inflammation is tissue infiltration by immune cells. Using the bulk RNA-seq data we also investigated the proportions of various immune cell types in relation to disease severity. First, we performed a candidate gene-based analysis and chose several immune marker genes to assess infiltration of T cells, T helper 1 (Th1) cells, T helper 2 (Th2) cells, B cells, basophils, neutrophils, eosinophils, and macrophages (Figure 2A). In contrast to *HK2* and the epithelial marker genes, the expression of these individual immune cell genes increased with the inflammation score. However, slight alterations in the expression patterns could be observed, mostly depending on the immune cell type. Expression of the T cell marker genes *TRAC*, *CD3D* and *CD3E,* the B cell marker gene *CD19,* the eosinophil marker gene *CCR2*, the neutrophil marker genes *CD14* and *CXCR4* as well as the basophil marker genes *ITGA2B* and *ITGA4* all gradually increased in a linear fashion with the inflammation score (Figure 2). In contrast, the marker genes for Th2 (*IL4*, *IL5*, *IL13*), Th1 (*IFNG*, *IL12*) cells and macrophages (*ITGAM*, *IL1B*) all only displayed biphasic expression patterns with a first phase characterized by slight increases in expression until an inflammation score of approximately 0.5 and a second phase at higher disease severity characterized by larger changes in their expression. The greatest expression changes were detected for the macrophage marker genes *ITGAM* and *IL1B* suggesting the greatest relative increase of these cells with increasing disease activity. Normalizing *HK2* expression to the candidate immune cell genes revealed gradual declines with increasing inflammation scores for all tested immune cell types indicating that the immune cell infiltration leads to more cells present in the mucosa, which do not or only express very little *HK2* (Figure 2B).

**Figure 2:**
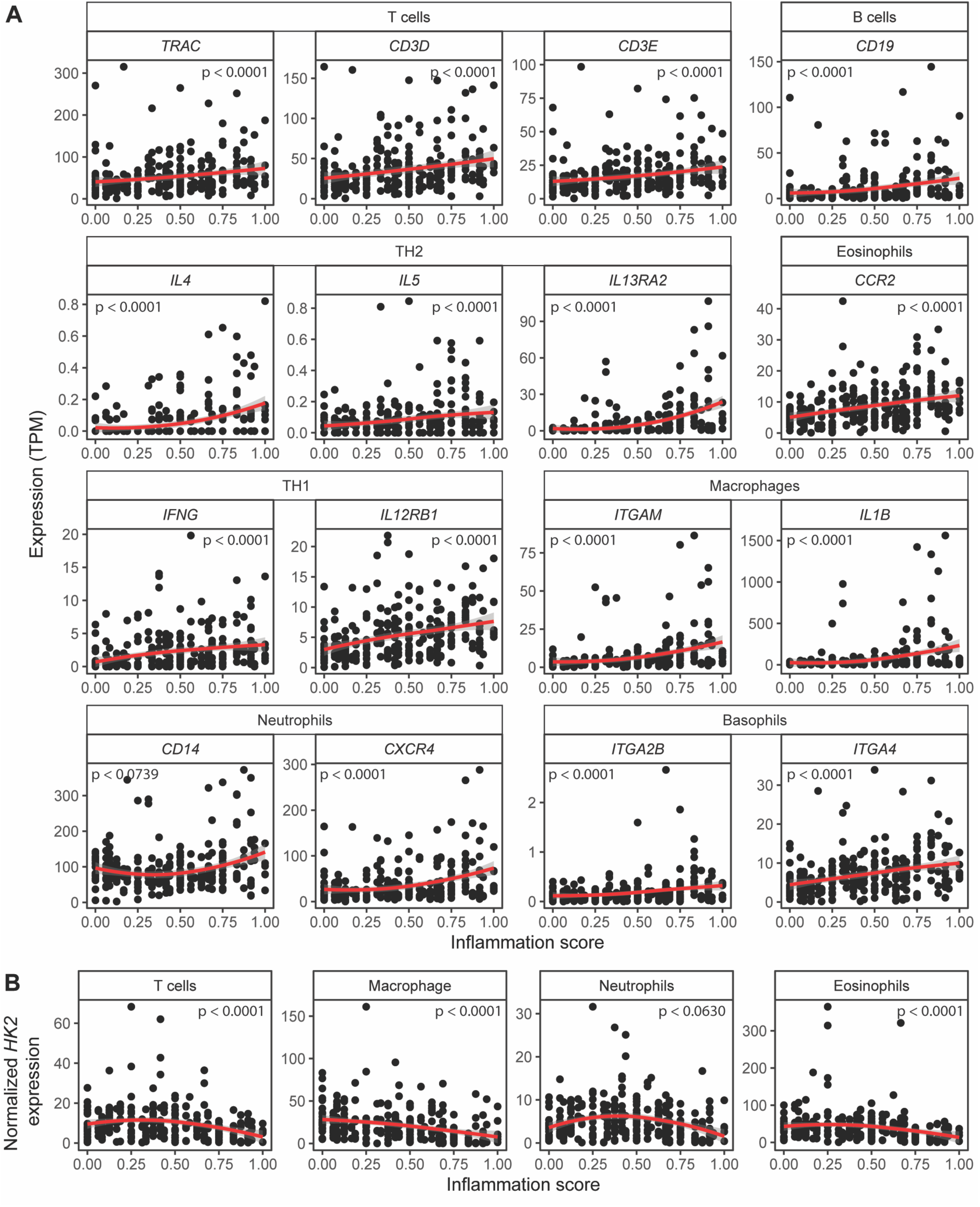
Expression of candidate immune marker genes increases with inflammation severity. Expression of selected marker genes, which are characteristic for individual immune cell types, were evaluated in patient biopsies. Expression of virtually all candidate immune marker genes increase with inflammation severity indicating an expansion of immune cells – a known feature of intestinal inflammation.

Next, we moved from the candidate gene-based to a systematic cellular deconvolution of the bulk RNA-seq data using MuSiC^10^. This program uses a reference single cell data set and cell type-specific expression profiles to derive the abundance of the individual cell types from bulk RNA-seq data. We restricted this analysis to the eight most abundant cell types. The proportion of epithelial cells gradually declined with increasing inflammation scores whereas the proportion of macrophages, B cells and regulatory T cells increased (Figure 3A). The proportion of mast, cytotoxic T cells and dendritic cells remained mostly unchanged. Therefore, this data indicates a massive gradual cellular remodeling of the intestinal mucosa during inflammation and the findings of the systematic cellular deconvolution supported those of the candidate gene-based analyses. Furthermore, the drastic decrease in the proportion of epithelial cells with increasing inflammation scores demonstrates again that the main source of *HK2* expression is lost during the disease course. This is therefore also reflected by the outcome that after accounting for the abundance of epithelial cells the normalized epithelial *HK2* expression gradually increased with disease severity (Figure 3B).

**Figure 3:**
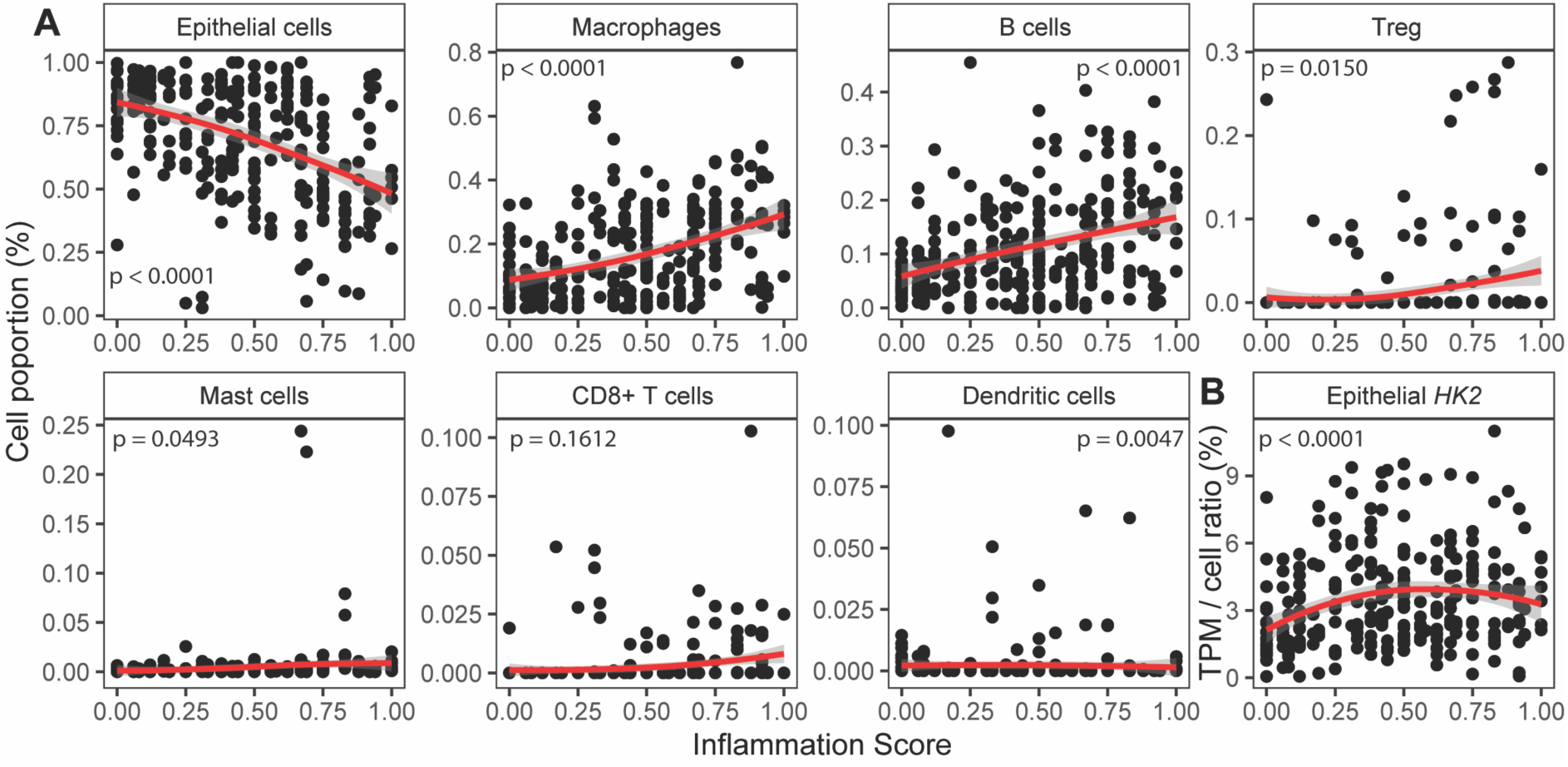
Deconvolution of mucosal transcriptomes demonstrates cellular remodeling during inflammation. **A)** MuSiC cell deconvolution package was used to estimate ratios of the most abundant cell types based on the mucosal patient transcriptomes. Notably, epithelial cell abundance is decreasing with inflammation, whereas abundances of macrophages and B cells, the two most frequent immune cell types that combine for approximately 35 % proportion, are increasing with inflammation severity. Therefore, cell deconvolution supports the candidate gene analyses indicating mucosal cellular remodeling characterized by a reduction of epithelial cells and an expansion of immune cells with increasing inflammation severity. **B)** *HK2* expression increases with inflammation scores after normalization to epithelial cell abundance.

### Apical HK2 expression in mature enterocytes correlates with inflammation severity

To validate the transcriptome data and to elucidate the biogeography of the changes in tissue architecture in relation to HK2 expression, we performed immunostainings on colonic biopsies of an independent set of UC patients that had been thoroughly scored for histological disease severity (Nancy score). HK2 protein levels increased non-significantly in mildly inflamed (score 1) compared to non-inflamed specimen (score 0) and remained unaltered at medium or strong inflammation (scores 2 and 4) (Figure 4A,C). In contrast, protein levels of the epithelial marker E-cadherin (ECAD) were unchanged in mild (score 1) but non-significantly reduced at medium or strong inflammation (scores 2 and 4) compared to non-inflamed control samples (Figure 4A,C). Notably, structural epithelial damage of inflamed biopsies was detectable with increasing inflammatory scores (Figure 4C) with loss of epithelial cells and an infiltration of other cells, probably immune cells, into the inflamed mucosa. In addition, many epithelial cells that were not shed but still present at the inflamed site had reduced ECAD expression, which could indicate transdifferention of these inflamed apical enterocytes. Normalizing the HK2 protein levels to those of ECAD revealed that epithelial HK2 protein levels (Figure 4B) significantly increased with inflammation regardless of disease severity (scores 1-4). This epithelial HK2 protein expression in inflamed intestinal mucosal biopsies mirrored and confirmed the transcript patterns of the RNA sequencing analysis.

**Figure 4:**
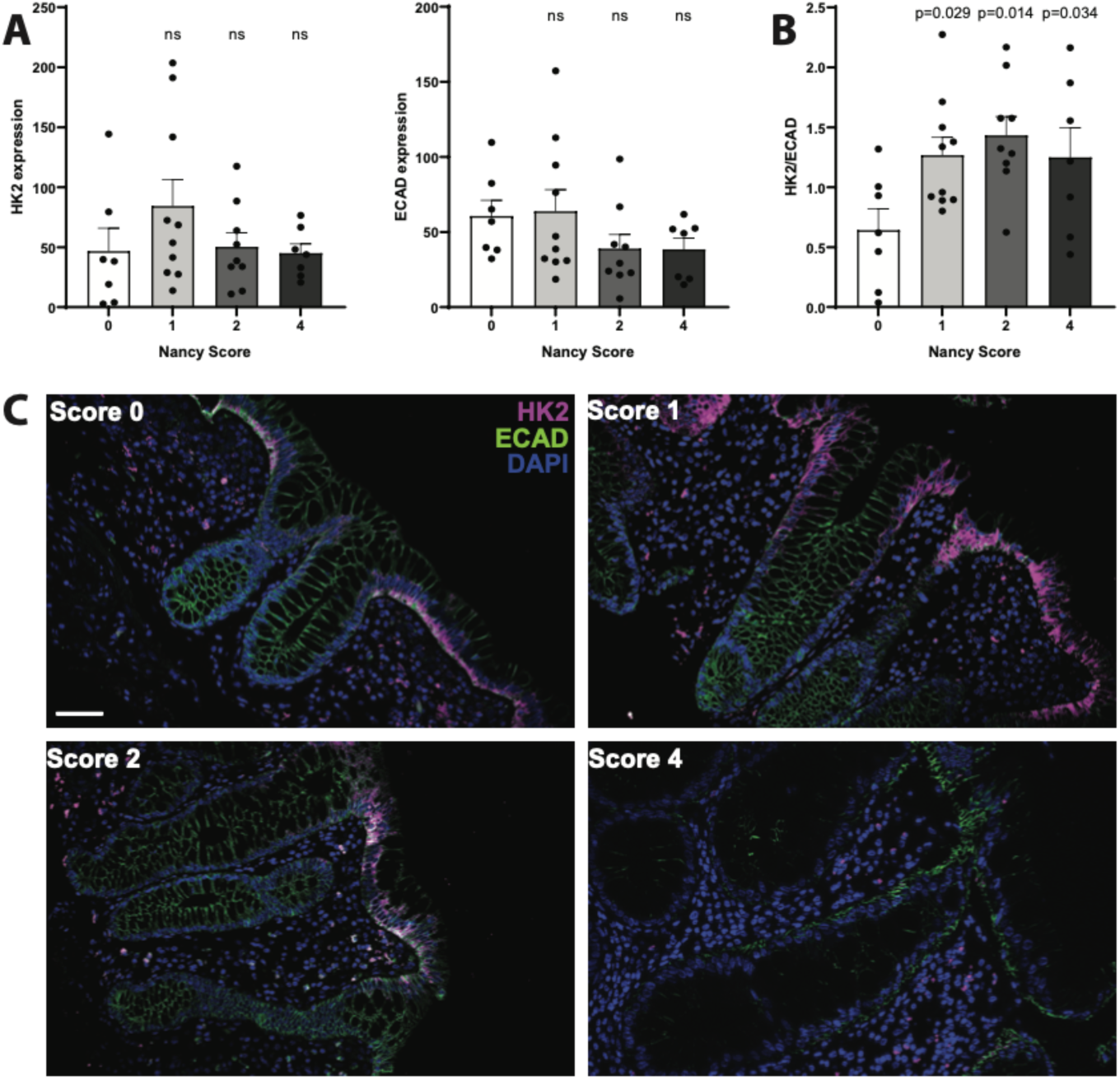
Epithelial HK2 protein expression increases with inflammation severity. Colonic mucosa biopsies of unrelated UC patients were stained for HK2, epithelial cells (ECAD) and nuclei (DAPI). All biopsies were analyzed for classical histological signs of inflammation using the Nancy Score (see method section). **A)** Quantification of HK2 and ECAD signal intensity in apical intestinal epithelium. Note that HK2 and ECAD level decrease with higher Nancy Scores. **B)** HK2 protein expression normalized to ECAD levels. Note that after epithelial normalization HK2 expression increases with inflammation severity, i.e. Nancy Score. Significance testing was performed using one-way ANOVA compared to Score = 0. **C)** Representative images of the multiplex immunofluorescence stainings (HK2, ECAD, DAPI). Scale bar indicates 50 µm. n = 7 – 10 per group. Note that microscopic images were taken from areas with rather intact epithelium for a better visualization of the mucosal structure.

Furthermore, we used recently-published data from single cell RNA sequencing of mucosal biopsies from IBD patients^11^ to investigate the *HK2* expression on a cellular level in relation to intestinal inflammation. This analysis of single cell RNA sequencing data confirmed our findings from the bulk RNA sequencing and immunofluorescence, as *HK2* was mainly expressed by mature enterocytes and then followed by goblet cells and immature enterocytes (Figure 5A,B). Next, we checked for *HK2* expression pattern in enterocytes at different inflammatory stages (healthy vs. inflamed vs. non-inflamed) and found that the lowest *HK2* expression can be observed in healthy controls, while cells from inflamed samples showed an increased HK2 expression (Figure 5C). Non-inflamed samples had intermediate *HK2* expression levels (Figure 5C), which indicates that HK2 expression is dysregulated in enterocytes of IBD patients also in the absence of an overt inflammation. This pattern of *HK2* expression was also present although less predominant in immature enterocytes and goblet cells. Finally, we looked into the number of these three cell types that are detected in the three health states and found that mature and immature enterocytes are lost during inflammation, while incomplete recovery can be observed in non-inflamed tissue of IBD patients (Figure 5D). Altogether, the single cell data therefore supports a cellular reprogramming during inflammation with loss of mature enterocytes (the main cell type of the apical epithelium) and a dysregulated *HK2* expression in inflamed apical epithelial cells.

**Figure 5:**
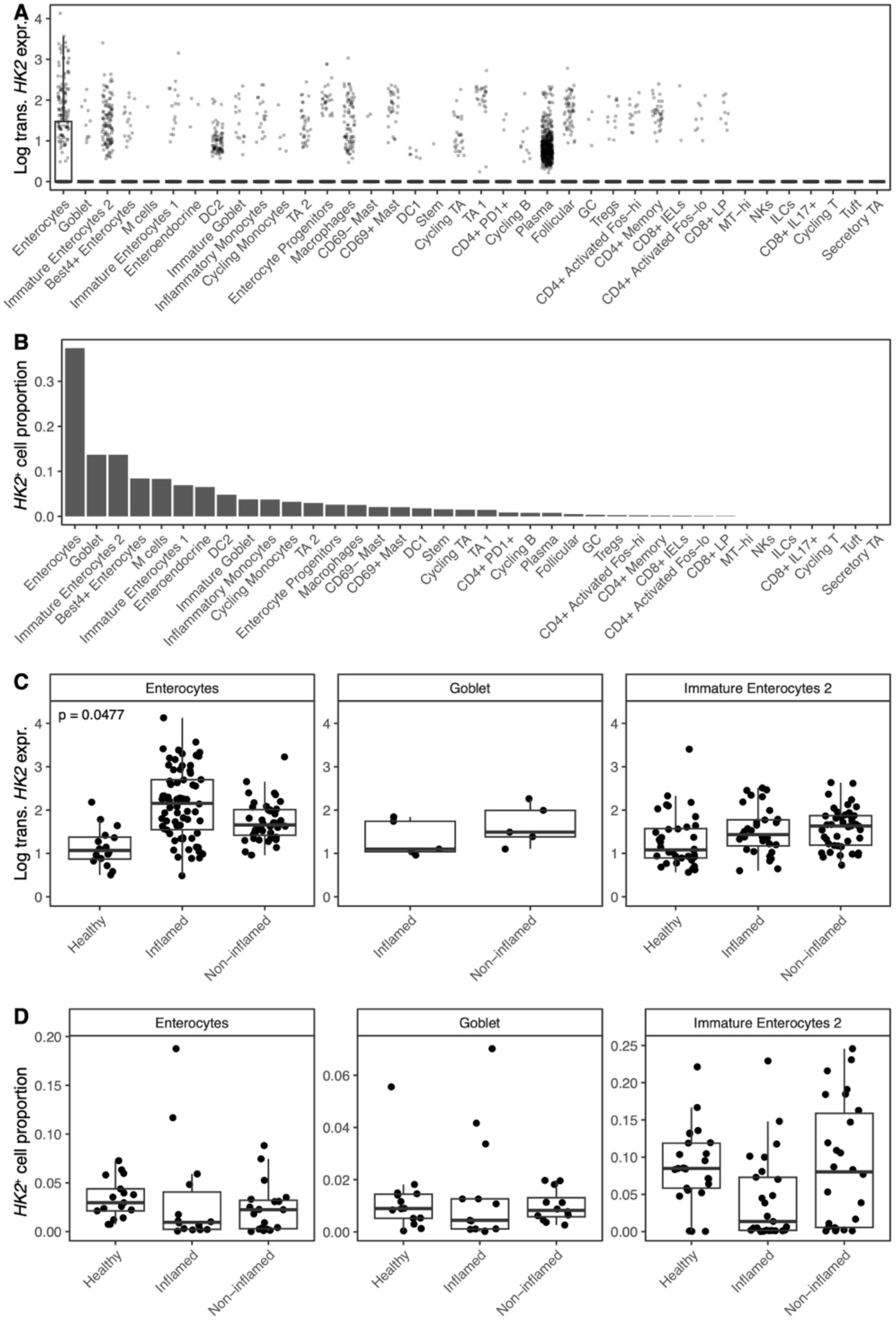
*HK2* is mainly expressed by mature enterocytes and increased during inflammation. *HK2* expression was analyzed in published single cell RNA sequencing data derived from mucosal biopsies of UC patients^11^. **A)** *HK2* expression per cell type. **B)** Abundances of cell types expressing *HK2*. **C)** *HK2* expression in mature & immature enterocytes and goblet cells divided into samples from healthy controls, inflamed and non-inflamed mucosa. **D)** Abundances of cell types expressing *HK2* after stratification into disease group.

In summary, both the RNA and protein data clearly demonstrate that epithelial HK2 was overexpressed during intestinal inflammation, which highlights HK2 as an indicator of active disease. Building upon previous findings from murine models that inhibition of HK2 expression either genetically or via the microbial metabolite butyrate protect from experimental colitis^6^, these new data now show that indeed HK2 expression is dysregulated in the mucosa of patients with active inflammation and therefore suggest that targeting HK2 may represent a promising approach to suppress intestinal inflammation in humans.

## Discussion

There is a growing need for valid disease biomarkers and molecular targets for intestinal inflammation as prevalence is increasing and despite significant efforts the cause of these diseases remains unknown. In a recent study, HK2 was identified as upregulated under inflammatory conditions in mice and humans irrespective of disease subtype (CD, UC or non-IBD colitis) and suppressing HK2 expression even protected from experimental colitis^6^ placing HK2 as a potential disease marker. However, we found other transcriptome datasets generated from severely inflamed patients in which HK2 expression was unaltered or even reduced^7^. Here, we therefore thoroughly investigated the relationship between HK2 expression and disease severity by analyzing transcriptome data derived from intestinal biopsies of patients with intestinal inflammation for which corresponding data on disease severity (HBI, Mayo score, Nancy score) were available. In addition, we performed cellular deconvolution analyses of the RNA sequencing data to infer the changes in cell proportions during inflammation. Finally, we used immunofluorescence staining to clarify the changes in HK2 protein biogeography in the mucosa during inflammation. Using these approaches, we demonstrated that raw HK2 RNA and protein expression at first gradually increased with the inflammation scores, yet after reaching a critical inflammation score HK2 expression declined again at very severe inflammation. These findings integrate the seemingly conflicting data from previous studies^7,12–14^ by linking HK2 expression to disease severity. We further were able to demonstrate that loss of HK2 expression at high inflammation levels was dependent on the disruption of brush border enterocytes as both the expression of epithelial marker genes were decreasing with the inflammation score and immunofluorescence analyses clearly pointed to a destruction of the apical epithelium – the main site of HK2 expression^15,16^. These findings are in line with clinical practice, in which epithelial damage is a key criterion to classify an increasing inflammatory state during IBD^17^, for example with epithelial erosion being a feature of histological inflammation in the Nancy score. By analyzing single cell RNA sequencing data derived from an independent set of UC patients, we confirmed that in the gut epithelial cells are the main source of *HK2* expression, both in terms of per cell expression as well as the number of cells contributing to overall expression (Figure 5). These cells are consequently lost due to increasing inflammation in the single cell data set, which would explain the decrease of *HK2* expression at high levels of inflammation^11^. Finally, we were able to deconstruct the simultaneous epithelial erosion and immune cell infiltration into the submucosa during intestinal inflammation from the RNA sequencing data. Based on the overall abundance and their induction (fold change) during inflammation, infiltrating macrophages seemed most relevant, but also other cell types such as B cells increased in proportion. Especially the expansion of macrophages could be important and contributing to the loss of epithelial cells and therefore HK2 expression during inflammation as IL-1β interferes with the tight junction complexes between IECs^11–13^ and thereby increases intestinal permeability^21^. Similar to epithelial erosion, immune cell infiltration is a classical feature of inflammation and used in clinical practice to histologically evaluate disease severity, in particular as a feature of the Nancy score^22^. In addition, transdifferentiation of HK2-positive mature enterocytes into other cell types such as HK2-negative immature enterocytes and stem cells^23,24^ or maybe even into mesenchymal cells as during epithelial– mesenchymal transition ^25^ could also contribute to a reduction of HK2 levels in the inflamed mucosa. In summary, our findings imply that during the course of inflammation, changes in the cellular composition of the mucosa, i.e. loss of brush border epithelial cells expressing high levels of HK2 and a corresponding massive recruitment of immune cells expressing little to no HK2 to the site of inflammation, affect the overall bulk HK2 expression leading to an overall reduction despite higher expression levels in fewer and fewer HK2-positive cells (Figure 6). Overall, we want to highlight the importance of taking into consideration the biogeography and changes in cell type expression ratios, when trying to identify disease biomarkers, especially regarding complex diseases such as intestinal inflammation involving various internal and external factors. Most previous studies used bulk expression data in the search for disease biomarkers or therapeutic targets and therefore may have missed other locally restricted but disease relevant genes like *HK2*. In our analyses HK2 expression was only significantly associated with disease severity after considering epithelial cell abundance (Figure 1E and Figure 3B) and we speculate that this pattern will also hold true for other genes with an apical or otherwise restricted expression. For example, other hexokinases (HK1, HKDC1), proteins responsible for the uptake of dietary nutrients (e.g. SGLT1, SLC15A1) or those contributing to the intestinal epithelial barrier (e.g. CLDN15, ZO-1) also show a predominant or even restricted apical expression and therefore possibly have not yet been identified as disease biomarker during severe intestinal inflammation due to the loss of their primary expression site. Notably, epithelial remodeling not only includes the partial loss of apical enterocytes and infiltration of lymphocytes, but also includes hyperregeneration, metaplasia and a loss of goblet cells. The advent of single cell technologies such as single cell RNA sequencing promise to pave the way for more refined analyses that will enable the discovery of more suitable disease markers and to enhance our understanding of the molecular and cellular mechanisms during disease progression.

**Figure 6:**
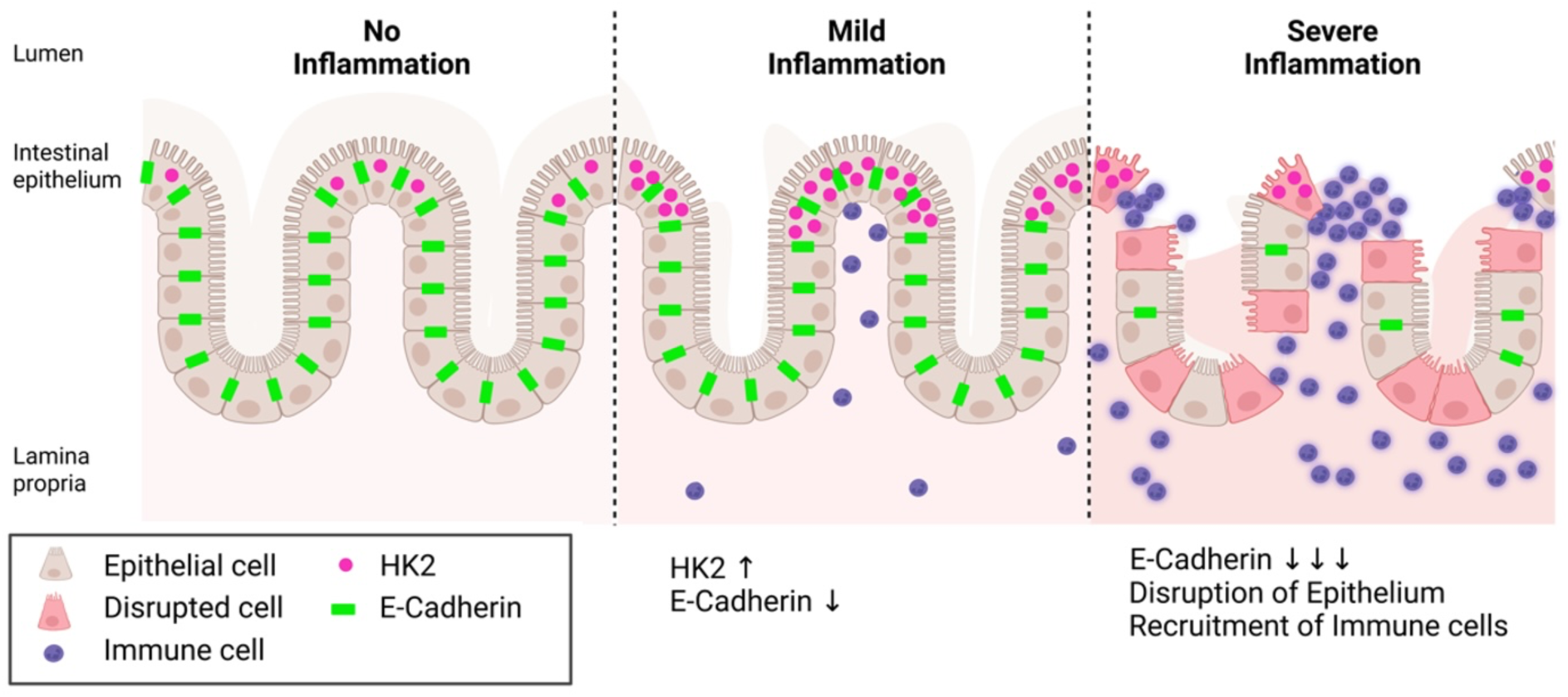
Model of HK2 expression changes in relation to intestinal inflammation severity. HK2 is predominantly expressed by apical epithelial cells and HK2 expression in these cells increases with disease severity. However, during the course of inflammation the cellular composition of the mucosa changes. In severely inflamed tissue, the epithelium is disrupted, brush border epithelial cells with high levels of HK2 are shed and lost, whereas immune cells with little to no HK2 expression are recruited. Therefore, overall HK2 expression is reduced in bulk tissue samples under severe inflammation despite local high expression in remaining HK2-positive epithelial cells. Created with BioRender.com

## Conclusions

HK2 has not yet been described as IBD risk gene, most probably due to its site-restricted expression, but here we were able to show an association of HK2 and epithelial status during intestinal inflammation. Therefore, epithelial HK2 expression correlates with disease severity making it a useful indicator of intestinal inflammation and highlighting the therapeutic potential of targeting HK2. However ultimately, clinical studies are required to indeed demonstrate the feasibility and efficacy of a HK2-targeted intervention in IBD patients.

## Declarations

### Ethics approval and consent to participate

All research complied with relevant ethical regulations, and usage of human tissue material and transcriptome data was approved by the ethics committee of the Medical Faculty at Kiel University (A156/03, A124/14, A102/16).

### Consent for publication

Not applicable

### Availability of data and material

Not applicable. All data was retrieved (see method section).

### Authors’ contributions

SW, LJ and FS designed research. SW, JT, LJ and FS performed experiments and analyzed the data. SS, CR, KA and PR provided samples and RNAseq data. SW, JT, LJ and FS prepared the figures. KA, PR, CK and FS obtained funding. SW, JT, LJ and FS co-wrote the manuscript with critical input from all authors. All authors read and approved the final manuscript.

## Acknowledgements

The authors thank Sabine Kock, Stefanie Baumgarten, Maren Reffelmann, Vivian Wegner, Tanja Klostermeier, Dorina Ölsner, Meike Hansen, Ronja Möhring and Sophie Reiher for excellent technical assistance. We further thank Florian Tran and Finn Hinrichsen for valuable input regarding the description of the clinical cohort characteristics.

## Authors’ information (optional)

Not applicable

---

**Figure S1:**
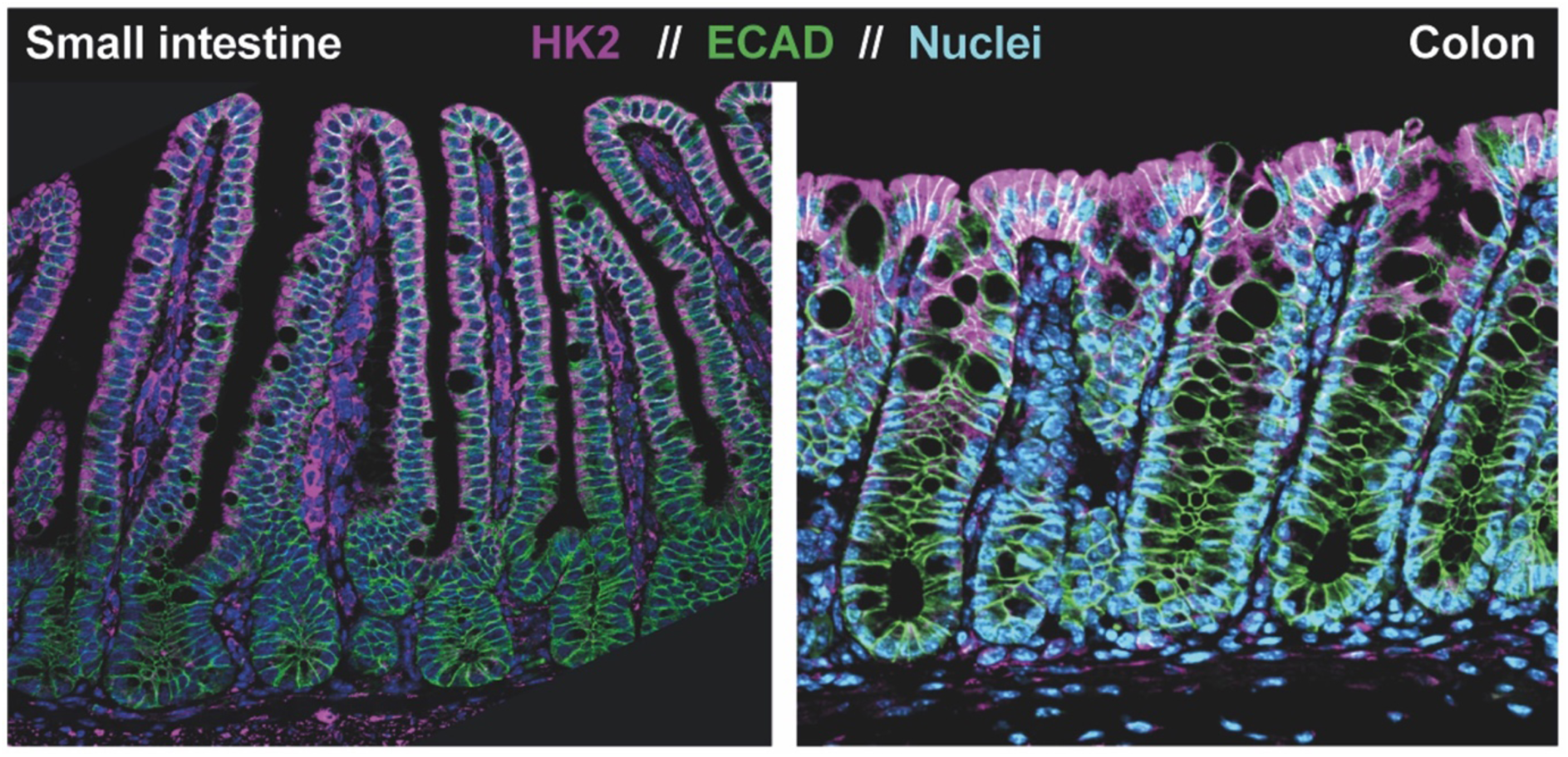
HK2 expression in the intestinal mucosa. Biopsies from small and large intestine were immuno-stained to localize the HK2 protein in the mucosa. Note that HK2 expression is mainly confined to epithelial cells of the apical mucosa both in the small and large intestine.

**Figure S2:**
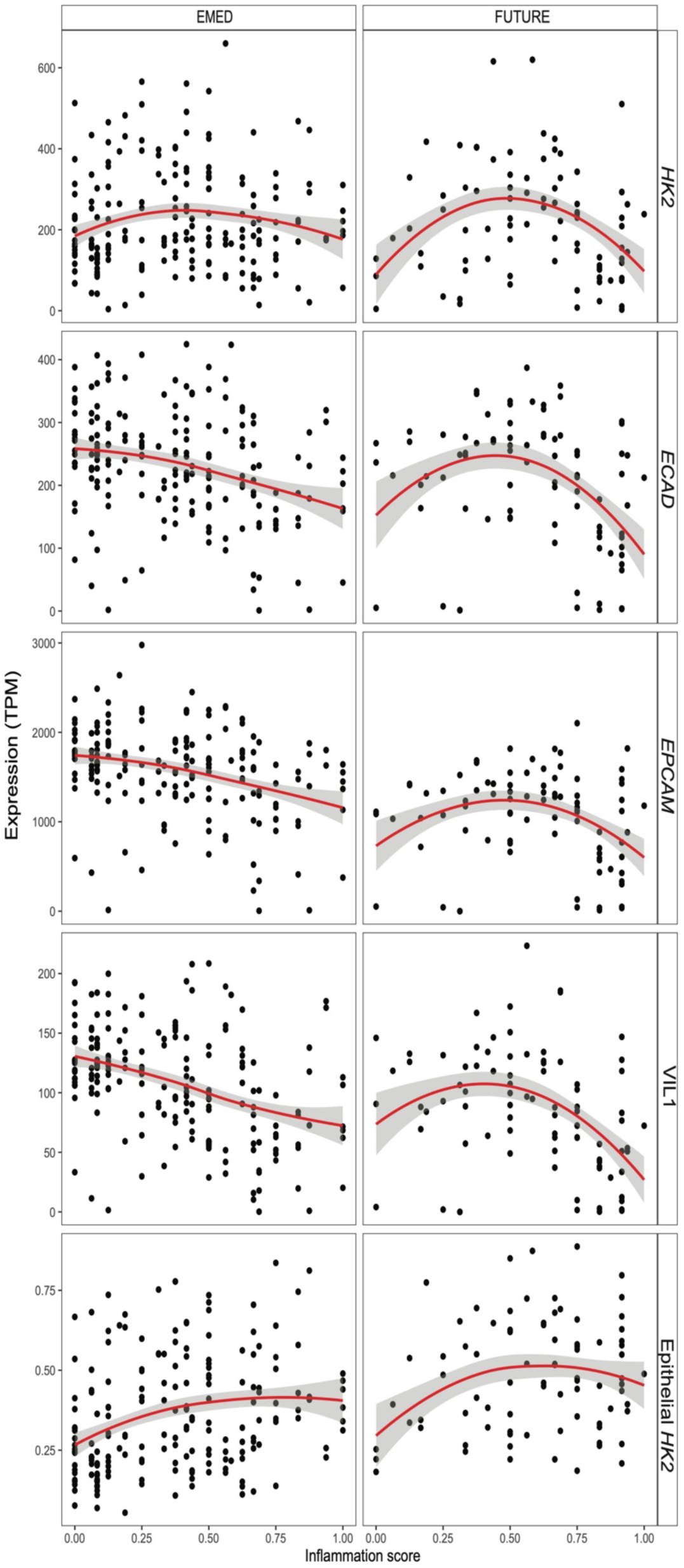
Epithelial *HK2* expression correlates with disease severity in both clinical cohorts. Expression data (TPM, transcript per million) of *HK2* and the epithelial marker genes *ECAD, VIL1* and *EPCAM* in the intestinal mucosa of patients with various degrees of gut inflammation split per clinical cohort FUTURE and EMED.

**Figure S3:**
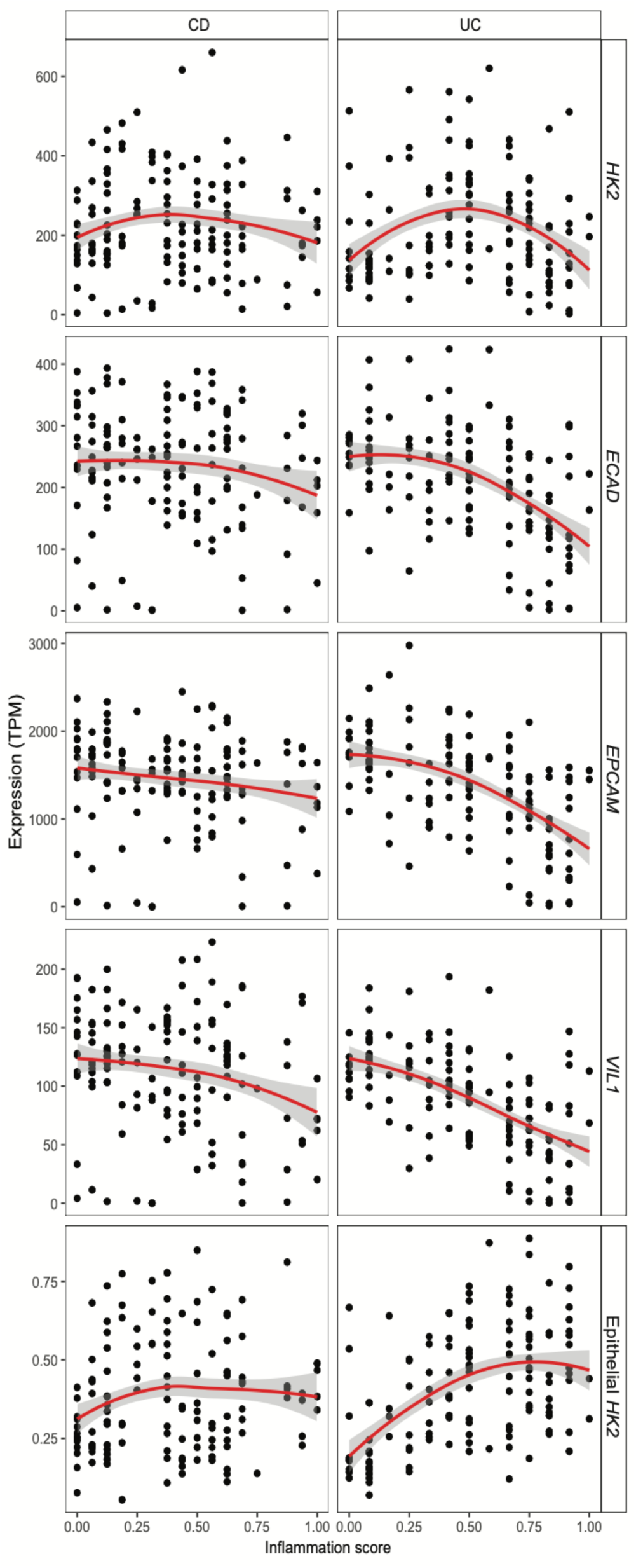
Epithelial *HK2* expression correlates with inflammation severity regardless of disease subtype. Expression data (TPM, transcript per million) of *HK2* and the epithelial marker genes *ECAD, VIL1* and *EPCAM* in the intestinal mucosa of patients with various degrees of gut inflammation split per disease subtype CD and UC.

**Figure S4:**
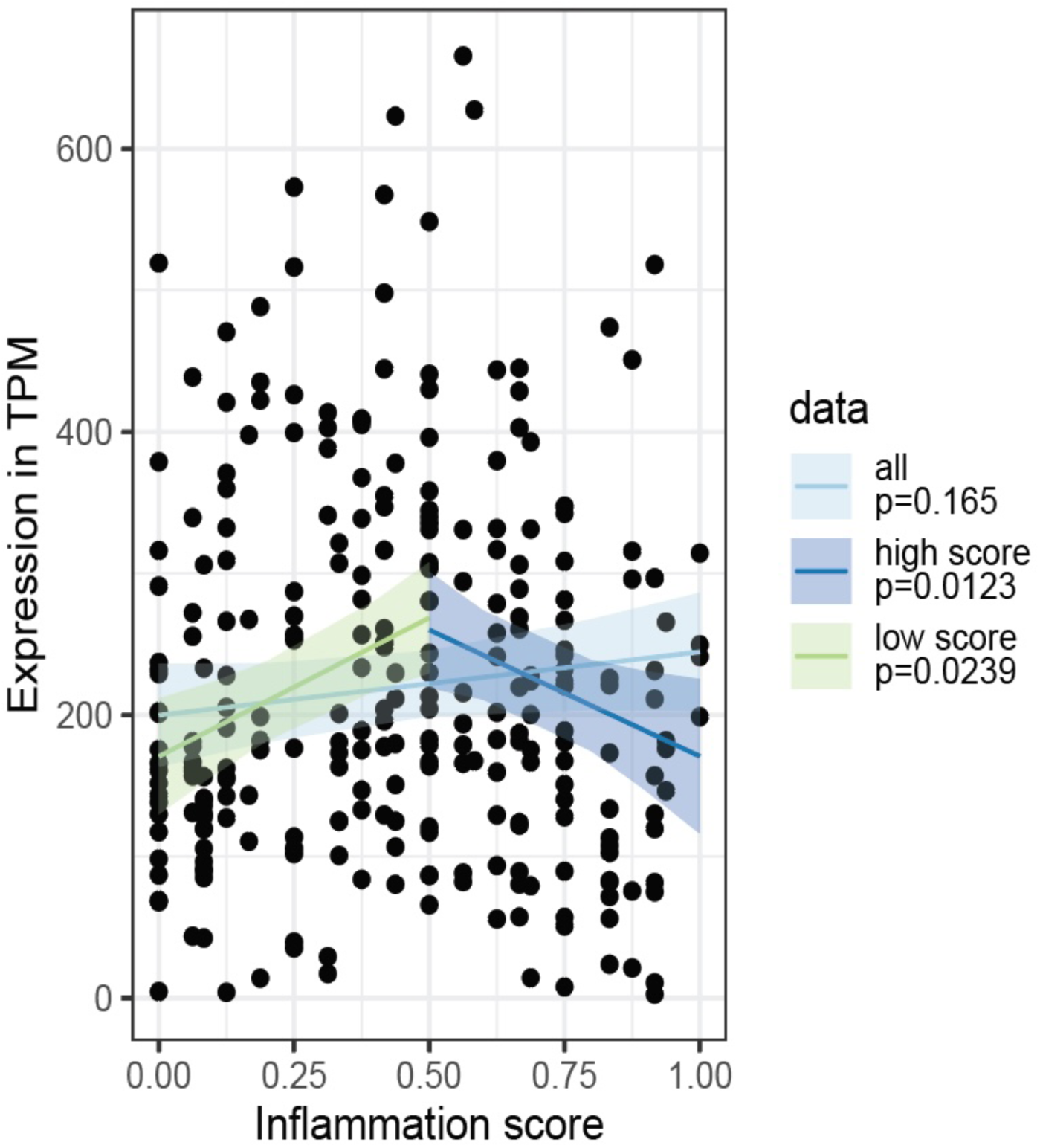
Correlation analysis for HK2 expression and inflammation score. Three different linear mixed effect models (PatientID as random factor) were fitted to the data. First, all data points were used to estimate the overall effect of *HK2* expression and inflammation score. Afterwards the data set was split into “low scores” (inflammation score < 0.5), “high scores” (inflammation score >0.5) and fitted models for these data subsets. While the first model showed no significant association between *HK2* expression and inflammation, there is a positive association at “low scores” and a negative association at “high scores” (see also table S2) indicating that *HK2* expression increases with inflammation during lower disease scores and then decreases at higher disease scores.

**Table S1:**
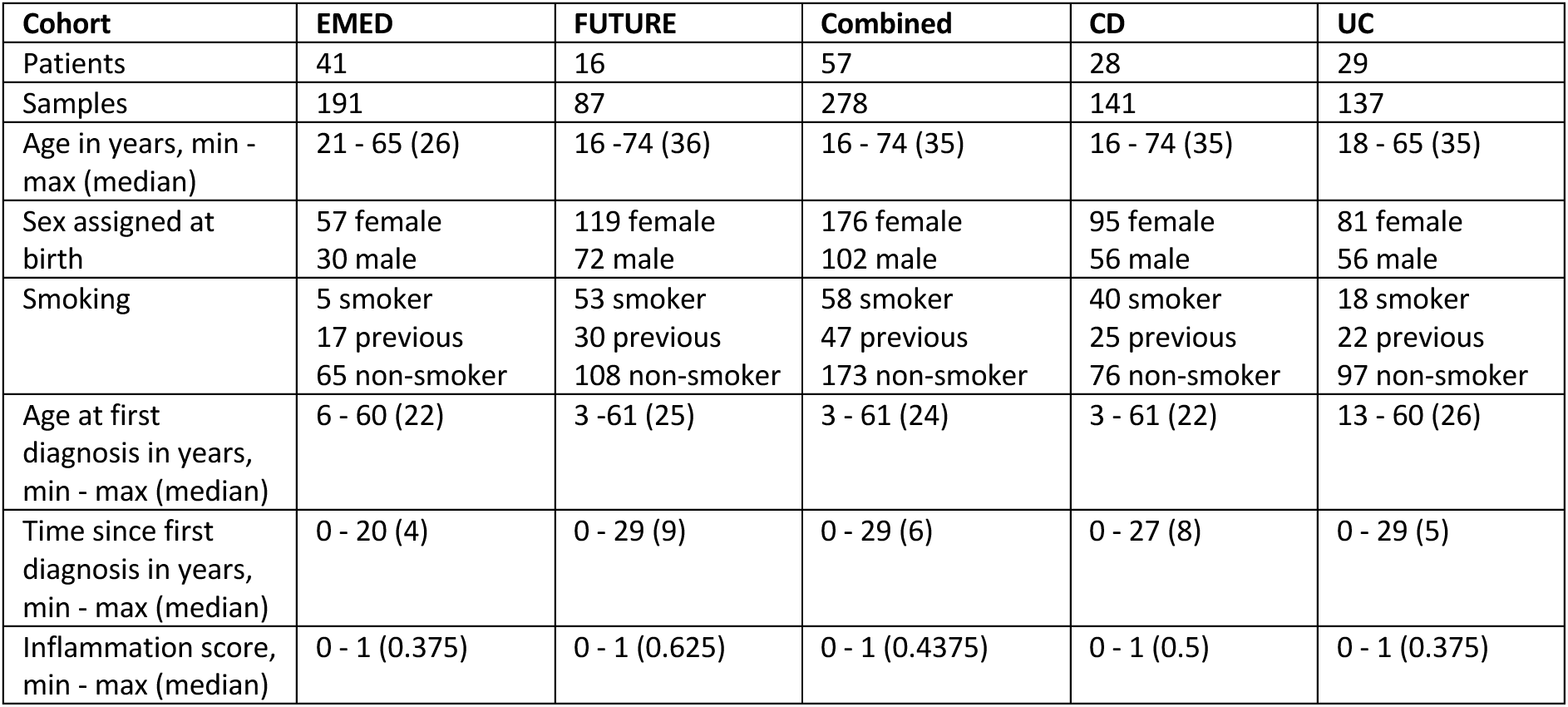
Cohort clinical characteristics.

**Table S2:** Comparison of linear mixed model statistics for the coefficients explaining HK2 expression with inflammation scores for models created with all data, low (score <0.5) or high (score >0.05) inflammation scores only.

|  | Estimate | Std. Error | DF | t-value | p-value |  |
| --- | --- | --- | --- | --- | --- | --- |
| **All data** | 44.73 | 32.07 | 179.94 | 1.395 | 0.165 |  |
| **Low score** | 195.86 | 59.48 | 154.44 | 3.293 | 0.0123 | ** |
| **High score** | -178.6 | 78 | 114.87 | -2.29 | 0.0239 | * |

**Table S3:** Translation of the HBI and Mayo Score into a general Inflammation Score. The definitions of the HBI and Mayo scores are listed. The Inflammation Score was calculated by scaling each the HBI and Mayo Score from 0-1 and then merging the scores.

| HBI |  |  |
| --- | --- | --- |
| Criterion | Score |  |
| General well-being | Very well | 0 |
|  | Slightly poor | 1 |
|  | Poor | 2 |
|  | Very poor | 3 |
|  | Terrible | 4 |
| Abdominal pain | None | 0 |
|  | Mild | 1 |
|  | Moderate | 2 |
|  | Severe | 3 |
| Abdominal mass | None | 0 |
|  | Dubious | 1 |
|  | Definite | 2 |
|  | Definite with tenderness | 3 |
| Stool frequency | Number of liquid stools | 1 for each liquid stool per day |
| Complications | Arthralgia, Uveitis, Erythema nodosum, Aphthous ulcers, Pyoderma gangrenosum, Anal fissure, New fistula, Abscess | 1 point each |

| Mayo Score |  |  |
| --- | --- | --- |
| Criterion | Score |  |
| Number of stools per day | Normal for the patient | 0 |
|  | 1-2 stools more than normal | 1 |
|  | 3-4 stools more than normal | 2 |
|  | 5+ stools more than normal | 3 |
| Rectal blood | No blood | 0 |
|  | Blood streaks in less the half of evacuations | 1 |
|  | Evidence of fresh blood in most of the evacuations | 2 |
|  | Bowel movements with fresh blood | 3 |
| Endoscopy | Normal or inactive disease | 0 |
|  | Mild disease (erythema, decreased vascular pattern, mild friability) | 1 |
|  | Moderate disease (marked erythema, lack of vascular pattern, friability, erosions) | 2 |
|  | Severe disease (spontaneous bleeding, ulceration) | 3 |
| Global medical assessment | Normal | 0 |
|  | Mild disease | 1 |
|  | Moderate disease | 2 |
|  | Severe disease | 3 |

| Calculation of inflammation score |  |  |  |  |  |  |  |  |
| --- | --- | --- | --- | --- | --- | --- | --- | --- |
| HBI | InfScore |  | Mayo | InfScore | >>> | InfScore | HBI | Mayo |
| 0 | 0.00 |  | 0 | 0.00 | merged | 0.00 | 0 | 0 |
| 1 | 0.06 |  | 1 | 0.08 |  | 0.06 | 1 |  |
| 2 | 0.12 |  | 2 | 0.17 |  | 0.08 |  | 1 |
| 3 | 0.18 |  | 3 | 0.25 |  | 0.12 | 2 |  |
| 4 | 0.24 |  | 4 | 0.33 |  | 0.17 |  | 2 |
| 5 | 0.29 |  | 5 | 0.42 |  | 0.18 | 3 |  |
| 6 | 0.35 |  | 6 | 0.50 |  | 0.24 | 4 |  |
| 7 | 0.41 |  | 7 | 0.58 |  | 0.25 |  | 3 |
| 8 | 0.47 |  | 8 | 0.67 |  | 0.29 | 5 |  |
| 9 | 0.53 |  | 9 | 0.75 |  | 0.33 |  | 4 |
| 10 | 0.59 |  | 10 | 0.83 |  | 0.35 | 6 |  |
| 11 | 0.65 |  | 11 | 0.92 |  | 0.41 | 7 |  |
| 12 | 0.71 |  | 12 | 1.00 |  | 0.42 |  | 5 |
| 13 | 0.76 |  |  |  |  | 0.47 | 8 |  |
| 14 | 0.82 |  |  |  |  | 0.50 |  | 6 |
| 15 | 0.88 |  |  |  |  | 0.53 | 9 |  |
| 16 | 0.94 |  |  |  |  | 0.58 |  | 7 |
| 17 | 1.00 |  |  |  |  | 0.59 | 10 |  |
|  |  |  |  |  |  | 0.65 | 11 |  |
|  |  |  |  |  |  | 0.67 |  | 8 |
|  |  |  |  |  |  | 0.71 | 12 |  |
|  |  |  |  |  |  | 0.75 |  | 9 |
|  |  |  |  |  |  | 0.76 | 13 |  |
|  |  |  |  |  |  | 0.82 | 14 |  |
|  |  |  |  |  |  | 0.83 |  | 10 |
|  |  |  |  |  |  | 0.88 | 15 |  |
|  |  |  |  |  |  | 0.92 |  | 11 |
|  |  |  |  |  |  | 0.94 | 16 |  |
|  |  |  |  |  |  | 1.00 | 17 | 12 |

**Table S4:** Original and pooled deconvolution cell types. Originally classified cell types that are ontogenetically related, e.g. Paneth cells, enterocytes, goblet cells and stem cells, were pooled and classified into “epithelial cells” to reduce cell type diversity.

| Original cell type description | Grouped into cell type |
| --- | --- |
| TA 1 | epis |
| TA 2 | epis |
| Enterocyte Progenitors | epis |
| Immature Goblet | epis |
| Immature Enterocytes 1 | epis |
| Immature Enterocytes 2 | epis |
| Cycling TA | epis |

## References

1. Kaplan, G. G. & Windsor, J. W. The four epidemiological stages in the global evolution of inflammatory bowel disease. Nat Rev Gastroenterol Hepatol 18, 56–66 (2021).

2. Kaplan, G. G. The global burden of IBD: from 2015 to 2025. Nat Rev Gastroenterol Hepatol 12, 720–727 (2015).

3. Rath, E. & Haller, D. Intestinal epithelial cell metabolism at the interface of microbial dysbiosis and tissue injury. Mucosal Immunology 15, 595–604 (2022).

4. Gaber, T., Strehl, C. & Buttgereit, F. Metabolic regulation of inflammation. Nature Reviews Rheumatology 13, 267–279 (2017).

5. Soto-Heredero, G., Gómez de las Heras, M. M., Gabandé-Rodríguez, E., Oller, J. & Mittelbrunn, M. Glycolysis – a key player in the inflammatory response. FEBS J 287, 3350–3369 (2020).

6. Hinrichsen, F. et al. Microbial regulation of hexokinase 2 links mitochondrial metabolism and cell death in colitis. Cell Metab 33, 2355–2366.e8 (2021).

7. Peters, L. A. et al. A functional genomics predictive network model identifies regulators of inflammatory bowel disease. Nat Genet 49, 1437–1449 (2017).

8. Schreiber, S. et al. Therapeutic Interleukin-6 Trans-signaling Inhibition by Olamkicept (sgp130Fc) in Patients With Active Inflammatory Bowel Disease. Gastroenterology 160, 2354–2366.e11 (2021).

9. Zeissig, S. et al. Vedolizumab is associated with changes in innate rather than adaptive immunity in patients with inflammatory bowel disease. Gut 68, 25–39 (2019).

10. Wang, X., Park, J., Susztak, K., Zhang, N. R. & Li, M. Bulk tissue cell type deconvolution with multi-subject single-cell expression reference. Nat Commun 10, 380 (2019).

11. Smillie, C. S. et al. Intra- and Inter-cellular Rewiring of the Human Colon during Ulcerative Colitis. Cell 178, 714–730.e22 (2019).

12. Marigorta, U. M. et al. Transcriptional risk scores link GWAS to eQTLs and predict complications in Crohn’s disease. Nat Genet 49, 1517–1521 (2017).

13. Haberman, Y. et al. Pediatric Crohn disease patients exhibit specific ileal transcriptome and microbiome signature. J. Clin. Invest. 124, 3617–3633 (2014).

14. Quraishi, M. N. et al. A Pilot Integrative Analysis of Colonic Gene Expression, Gut Microbiota, and Immune Infiltration in Primary Sclerosing Cholangitis-Inflammatory Bowel Disease: Association of Disease With Bile Acid Pathways. Journal of Crohn’s and Colitis 14, 935–947 (2020).

15. Sommer, F., Nookaew, I., Sommer, N., Fogelstrand, P. & Bäckhed, F. Site-specific programming of the host epithelial transcriptome by the gut microbiota. Genome Biol 16, 62 (2015).

16. The Tabula Muris Consortium et al. Single-cell transcriptomics of 20 mouse organs creates a Tabula Muris. Nature 562, 367–372 (2018).

17. Guan, Q. A Comprehensive Review and Update on the Pathogenesis of Inflammatory Bowel Disease. Journal of Immunology Research 2019, 1–16 (2019).

18. Coccia, M. et al. IL-1β mediates chronic intestinal inflammation by promoting the accumulation of IL-17A secreting innate lymphoid cells and CD4+ Th17 cells. Journal of Experimental Medicine 209, 1595–1609 (2012).

19. Chotikatum, S., Naim, H. Y. & El-Najjar, N. Inflammation induced ER stress affects absorptive intestinal epithelial cells function and integrity. International Immunopharmacology 55, 336–344 (2018).

20. Lin, Y. et al. Non-hematopoietic STAT6 induces epithelial tight junction dysfunction and promotes intestinal inflammation and tumorigenesis. Mucosal Immunology 12, 1304–1315 (2019).

21. He, L. et al. Gut Epithelial Vitamin D Receptor Regulates Microbiota-Dependent Mucosal Inflammation by Suppressing Intestinal Epithelial Cell Apoptosis. Endocrinology 159, 967–979 (2018).

22. Marchal-Bressenot, A. et al. Development and validation of the Nancy histological index for UC. Gut 66, 43–49 (2017).

23. Moor, A. E. et al. Spatial Reconstruction of Single Enterocytes Uncovers Broad Zonation along the Intestinal Villus Axis. Cell 175, 1156–1167.e15 (2018).

24. Wei, X. et al. Extensive jejunal injury is repaired by migration and transdifferentiation of ileal enterocytes in zebrafish. Cell Reports 42, 112660 (2023).

25. López-Novoa, J. M. & Nieto, M. A. Inflammation and EMT: an alliance towards organ fibrosis and cancer progression. EMBO Mol Med 1, 303–314 (2009).

26. Harvey, R. F. & Bradshaw, J. M. A Simple Index Of Crohn’s-Disease Activity. The Lancet 315, 514 (1980).

27. R Core Team. R: A Language and Environment for Statistical Computing. (R Foundation for Statistical Computing, Vienna, 2022).

28. Bates, D., Mächler, M., Bolker, B. & Walker, S. Fitting Linear Mixed-Effects Models Using lme4. Journal of Statistical Software 67, 1–48 (2015).

29. Kuznetsova, A., Brockhoff, P. B. & Christensen, R. H. B. lmerTest Package: Tests in Linear Mixed Effects Models. Journal of Statistical Software 82, 1–26 (2017).

30. Fox, J. & Weisberg, S. An R Companion to Applied Regression. (Sage, Thousand Oaks CA, 2019).

31. Wickham, H. Ggplot2: Elegant Graphics for Data Analysis. (Springer-Verlag, New York, 2016).

32. Love, M. I., Huber, W. & Anders, S. Moderated estimation of fold change and dispersion for RNA-seq data with DESeq2. Genome Biol 15, 550 (2014).

33. Hao, Y. et al. Integrated analysis of multimodal single-cell data. Cell 184, 3573–3587.e29 (2021).

